# Population Genetics Based Phylogenetics Under Stabilizing Selection for an Optimal Amino Acid Sequence: A Nested Modeling Approach

**DOI:** 10.1101/120238

**Authors:** Jeremy M. Beaulieu, Brian C. O’Meara, Russell Zaretzki, Cedric Landerer, Juanjuan Chai, Michael A. Gilchrist

**Author notes:** 50 Main St, Suite 1039, White Plains, NY 10606. **Associate Editor:** TBD.

## Abstract

We present a new phylogenetic approach SelAC (Selection on Amino acids and Codons), whose substitution rates are based on a nested model linking protein expression to population genetics. Unlike simpler codon models which assume a single substitution matrix for all sites, our model more realistically represents the evolution of protein coding DNA under the assumption of consistent, stabilizing selection using cost-benefit approach. This cost-benefit approach allows us generate a set of 20 optimal amino acid specific matrix families using just a handful of parameters and naturally links the strength of stabilizing selection to protein synthesis levels, which we can estimate. Using a yeast dataset of 100 orthologs for 6 taxa, we find SelAC fits the data much better than popular models by 10^4^–10^5^ AICc units. Our results indicate there is great potential for more accurate inference of phylogenetic trees and branch lengths from already existing data through the use of nested, mechanistic models. Additional parameters estimated by SelAC indicate that a large amount of non-phylogenetic, but biologically meaningful, information can be inferred from exisiting data. For example, SelAC prediction of gene specific protein synthesis rates correlates well with both empirical (*r*=0.33−0.48) and other theoretical predictions (*r*=0.45−0.64) for multiple yeast species. SelAC also provides estimates of the optimal amino acid at each site. Finally, because SelAC is a nested approach based on clearly stated biological assumptions, future modifications, such as including shifts in the optimal amino acid sequence within or across lineages, are possible.

## Introduction

Phylogenetic analyses plays a critical role in most aspects of biology, particularly in the fields of ecology, evolution, paleontology, medicine, and conservation. While the scale and impact of phylogenetic studies have increased substantially over the past two decades, the realism of the mathematical models on which these analyses are based has changed relatively little by comparison. The most popular models of DNA substitution used in molecular phylogenetics are simple nucleotide models that date back the early 1980’s and 90’s, e.g. F81, F84, HYK85, TN93, and GTR (see Yang (2014) for an overview), and are indifferent to the type of sequences they are fitted to. For example, when evaluating protein-coding sequences these models are inherently agnostic with regards to the different amino acid substitutions and their impact on gene function and, as a result, cannot describe the behavior of natural selection at the amino acid or protein level.

Two important and independent attempts to address this critical shortcoming were introduced by Goldman and Yang (1994, commonly abbreviated as GY94) and Muse and Gaut (1994). These models were explicitly built for protein coding data, assuming that differences in the physicochemical properties between amino acids, or physicochemical distances for short, could affect substitution rates. These physicochemical based codon models as originally introduced have rarely been used for empirical data. Instead, these often cited models have served as the basis for an array of simpler and, in turn, more popular *ω* models that, starting with Nielsen and Yang (1998); Yang and Nielsen (1998), typically assume an equal fixation probability for *all* non-synonymous mutations. Although often attributed to GY94, these later and simpler models were the first to employ the single term *ω* to model the differences in fixation probability between nonsynonomous and synonomyous changes at all sites. Since their introduction, more complex models have been developed that allow *ω* to vary between sites or branches (as cited in Anisimova, 2012) and include selection on different synonyms for the same amino acid (e.g. Yang and Nielsen, 2008)

In Goldman and Yang (1994); Nielsen and Yang (1998); Yang and Nielsen (1998) and later studies based on their work, *ω* is suggested to indicate whether a given site within a protein sequence is under consistent ‘stabilizing (*ω*< 1) or ‘diversifying’ (*ω*> 1) selection. Contrary to popular belief, *ω* does not describe whether a site is evolving under a constant regime of stabilizing or diversifying selection, but instead how a very particular *selective environment* changes over time. Below we explain how the actual behavior of these models is inconsistent with how ‘stabilizing’ and ‘diversifying’ selection are otherwise defined and understood (e.g. see Pellmyr, 2002).

For example, when *ω*<1, synonymous substitutions have a higher substitution rate than any possible non-synonymous substitutions. As a result, the model behaves as if the resident amino acid *i* at a given site is favored by natural selection. Even when *ω* is allowed to vary between sites, symmetrical aspects of the model means that for any given site the strength of selection for the resident amino acid *i* over its 19 alternatives is equally strong regardless of their physicochemical properties. Paradoxically, natural selection for amino acid *i* persists *until* a substitution for another amino acid, *j*, occurs. As soon as amino acid *j* fixes, but not before, selection now favors amino acid *j* equally over all other amino acids, including amino acid i. This is now the opposite scenario from when *i* was the resident. Thus, the simplest and most consistent interpretation of *ω* is that it represents the rate at which the selective environment itself changes, and this change in selection perfectly coincides with the fixation of a new amino acid.

Similarly, when *ω*> 1, synonymous substitutions have a lower substitution rate than any possible non-synonymous substitutions from the resident amino acid. Again due to the model’s symmetrical nature, the selection *against* the resident amino acid *i* is equally strong relative to alternative amino acids. The selection against the resident amino acid *i* persists until a substitution occurs at which point selection now *favors* amino acid i, as well as the 19 other amino acids, to the same degree *i* was previously disfavored. Given this behavior, *ω* based models are likely to only reasonably approximate a subset of scenarios such as perfectly symmetrical over-/under-dominance or positive/negative frequency dependent selection (Hughes and Nei, 1988; Nowak, 2006). Further, *ω* based models implicitly assumes the substitution is on the same timescale as the shifts in the optimal (or pessimal) amino acid.

## New Approaches

To address these fundamental shortcomings in *ω* based phylogenetic approaches, we present an approach where selection explicitly favors minimizing the cost-benefit function *η* of a protein whose relative performance is determined by the order and physicochemical properties of its amino acids. Our approach, which we call Selection on Amino acids and Codons or SelAC, is developed in the same vein as previous phylogenetic applications of the Wright-Fisher process (e.g. Dimmic et al., 2000; Halpern and Bruno, 1998; Koshi and Goldstein, 1997; Koshi et al., 1999; Lartillot and Philippe, 2004; Muse and Gaut, 1994; Rodrigue and Lartillot, 2014; Rodrigue et al., 2005; Thorne et al., 2012; Yang and Nielsen, 2008). Similar to Lartillot and Philippe (2004) and Rodrigue and Lartillot (2014), we assume there is a finite set of rate matrices describing the substitution process and that each position within a protein is assigned to a particular rate matrix category. Unlike that work, we assume *a priori* there are 20 different families of rate matrices, one family for when a given amino acid is favored at a site. The key parameters underlying these matrices are shared across genes except for gene expression. As a result, SelAC identifies the amino acid at a particular position within a protein that is favored by natural selection using a simple cost-benefit approach.

While natural selection on protein coding regions can take many forms, one general approach to describing its effects is by relating a codon sequence to the cost of producing the encoded protein and the functional benefit (or potential harm) from translating its sequence. The gene specific cost of protein synthesis can be affected by the amino acids used, the direct and indirect costs of peptide assembly by the ribosome, and the use of chaperones to aid in folding. Importantly, these costs can be computed to varying degrees of realism (e.g. Lynch and Marinov, 2015; Wagner, 2005). We have previously presented models of protein synthesis costs that, alternatively, take into account the cost of ribosome pausing (Shah and Gilchrist, 2011) or premature termination errors (Gilchrist et al., 2009; Gilchrist, 2007; Gilchrist and Wagner, 2006).

Protein function or ‘benefit’ can be affected by the amino acids at each site and their interactions. Linking amino acid sequence to protein function is a daunting task; thus for simplicity, we assume that for any given desired biological function to be carried out by a protein, that (a) the biological importance of this protein function is invariant across the tree, (b) single optimal amino acid sequence that carries out this function best, and (c) the functionality of alternative amino acid sequences declines with their physicochemical distance from the optimum on a site by site basis.

Beyond fitting the phylogenetic data better according to model adequacy and AICc, SelAC also makes inferences about other important biological processes. By comparing these inferences to other empirical data, such as we do with protein synthesis data, we can evaluate SelAC’s performance independent of the data it is fitted to. Indeed, SelAC’s assumptions lead to mechanistic and, thus, testable hypothesis about the nature of and relationships between mutation, protein function, gene expression, and rates of evolution. More importantly, alternative hypotheses could be used in place of ours and, in turn, phylogenetic and other types of data could be used to evaluate the support of these alternative models. Our hope is that by moving away from the more phenomenological models we can better connect population genetics, molecular biology, and phylogenetics allowing each area inform the others more effectively.

## Results

By linking transition rates *q*_*i*,*j*_ to gene expression in the form of protein synthesis rate *ϕ*, our approach allows use of the same model for genes under varying degrees of stabilizing selection. Specifically, we assume the strength of stabilizing selection for the optimal sequence, 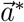, is proportional to the average protein synthesis rate *ϕ*, which we can estimate for each gene. In regards to model fit, our results clearly indicated that linking the strength of stabilizing selection for the optimal sequence to gene expression substantially improves our model fit. Further, including the shape parameter *α*_*G*_ for the random effects term *G*~Gamma(shape = *α*_*G*_,rate = *α*_*G*_) to allow for heterogeneity in this selection between sites within a gene improves the ΔAICc of SelAC+Γ over the simpler SelAC models by over 22,000 AIC units. Using either ΔAICc or AIC_w_ as our measure of model support, the SelAC models fit extraordinarily better than GTR + Γ, GY94, or FMutSel (Table 1). This is in spite of the need for estimating the optimal amino acid at each position in each protein, which accounts for 49,881 additional model parameters. Even when compared to the next most parameter rich codon model in our model set, FMutSel, SelAC+Γ model shows over 160,000 AIC unit improvement over FMutSel.

**Table 1.**
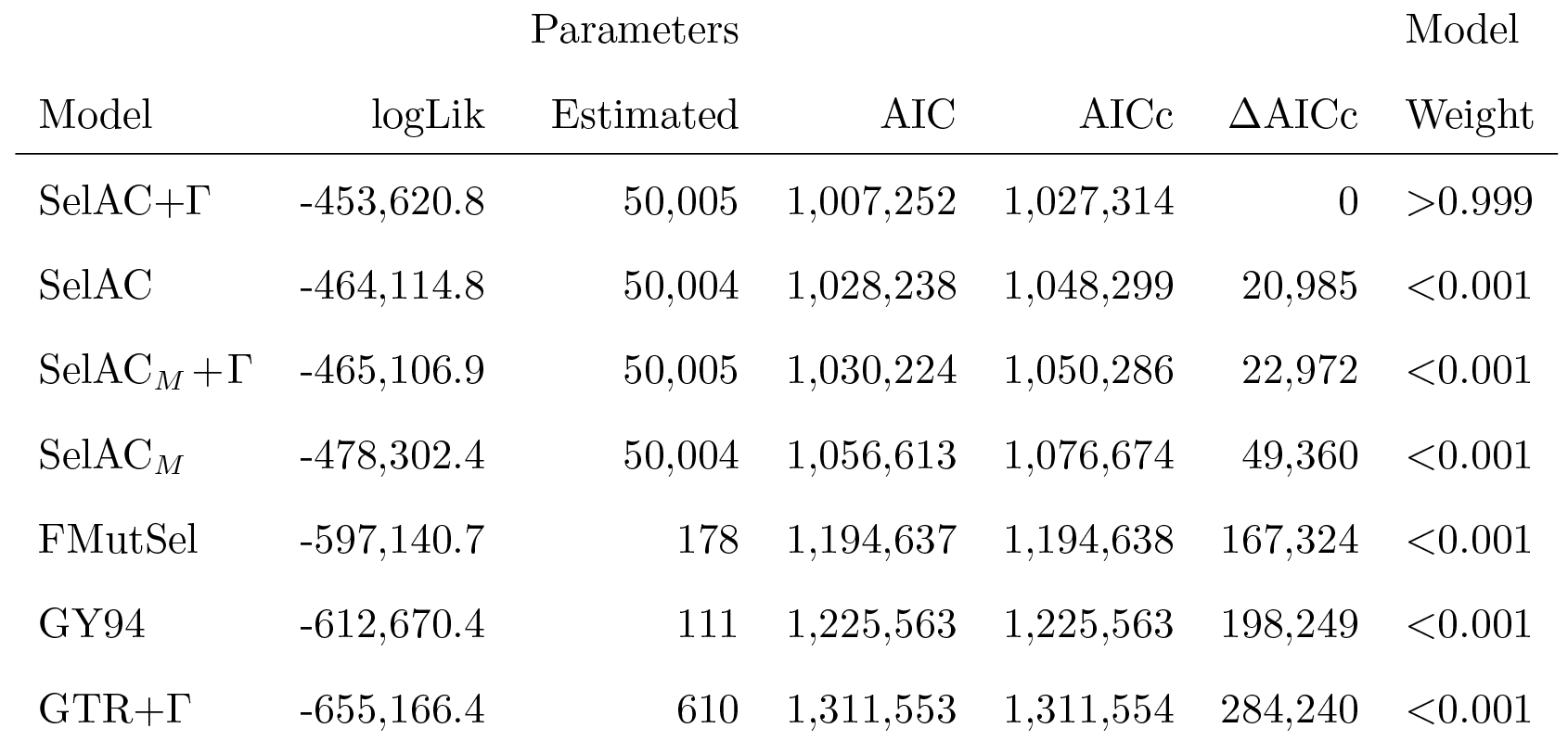
Comparison of model fits using AIC, AIC_c_, and AIC_w_. Note the subscripts *M* indicate model fits where the most common or ‘majority rule’ amino acid was fixed as the optimal amino acid *a*\* for each site. As discussed in text, despite the fact that *a*\* for each site was not fitted by our algorithm, its value was determined by examining the data and, as a result, represent an additional parameter estimated from the data and are accounted for in our table. Also, the sample size used in the calculation of AICc is assumed to be equal to the size of the matrix (number of taxa × number of sites).

The analysis building upon Jhwueng et al. (2014) suggests that using the number of taxa times the number of sites as the sample size performs best as a small sample size correction for estimating Kullback-Liebler distance in phylogenetic models (Appendix 1). This also has an intuitive appeal. In models that have at least some parameters shared across sites and some parameters shared across taxa, increasing the number of sites and/or taxa should be adding more samples for the parameters to estimate. This is consistent considering how likelihood is calculated for phylogenetic models: the likelihood for a given site is the sum of the probabilities of each observed state at each tip, which is then multiplied across sites. It is arguable that the conventional approach in comparative methods is calculating AICc in the same way. That is, if only one column of data (or “site”) is examined, as remains remarkably common in comparative methods, when we refer to sample size, it is technically the number of taxa multiplied by number of sites, even though it is referred to simply as the number of taxa.

With respect to estimates of *ϕ* within SelAC, they were strongly correlated with both empirical measurements (Pearson *r*=0.33−0.48) and theoretical predictions (Pearson *r*=0.45−0.64) of gene expression (Figure 1 and Figures S1-S2, respectively). In other words, using only codon sequences, our model can predict which genes have high or low expression levels. The estimate of the *α*_*G*_ parameter, which describes the site-specific variation in sensitivity of the protein’s functionality, indicated a moderate level of variation in gene expression among sites. Our estimate of *α*_*G*_ = 1.36, produced a distribution of sensitivity terms G ranged from 0.342-7.32, but with more than 90% of the weight for a given site-likelihood being contributed by the 0.342 and 1.50 rate categories. In simulation, however, of all the parameters in the model, only *α*_*G*_ showed a consistent bias, in that the MLE were generally lower than their actual values (see Supporting Materials). Other parameters in the model, such as the Grantham weights, provide an indication as to the physicochemical distance between amino acids. Our estimates of these weights only strongly deviate from Grantham’s 1974 original estimates in regards to composition weight, *α*_*c*_, which is the ratio of non-carbon atoms in the end groups or rings to the number of carbon atoms in side chains. Our estimate of the composition weighting factor of *α*_*c*_=0.459 is 1/4th the value estimate by Grantham which suggests that the substitution process is less sensitive to this physicochemical property when shared ancestry and variation in stabilizing selection are taken into account.

**FIG. 1.**
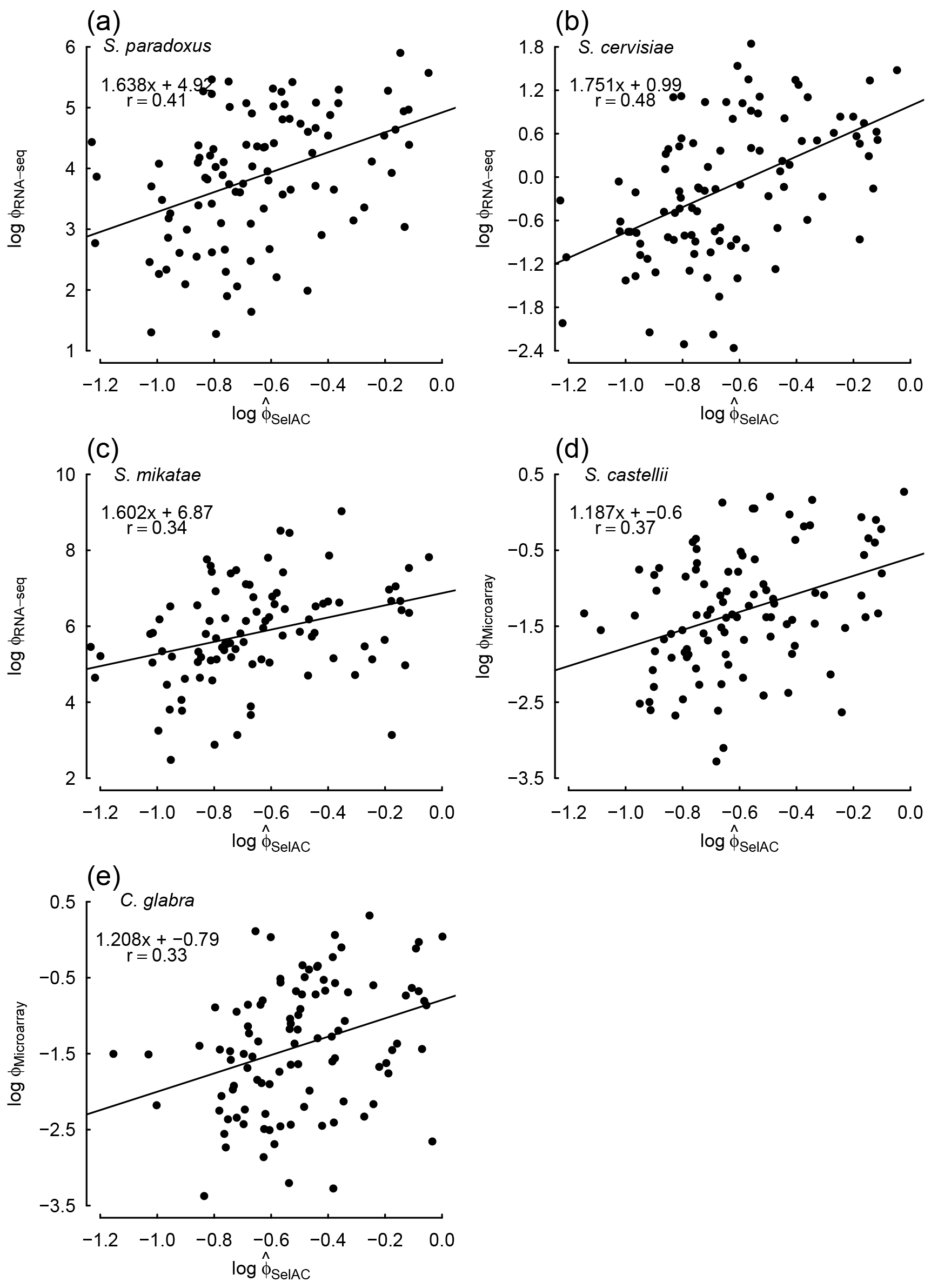
Comparisons between estimates of average protein translation rate 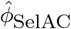 obtained from SelAC+Γ and direct measurements of expression for individual yeast taxa across the 100 selected genes from Salichos and Rokas (2013) measured during log-growth phase. Estimates of 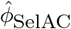 were generated by dividing the composite term *ψ*′ by 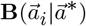. Gene expression was measured using either RNA-Seq (a)-(c) or microarray (d)-(e). The equations in the upper left hand corner of each panel represent the regression fit and the Pearson correlation coefficient *r*.

It is important to note that the nonsynonymous/synonymous mutation ratio, or *ω*, which we estimated for each gene under the FMutSel model strongly correlated with our estimates of *ϕ*′=*ψ*′/**B** where **B** depends on the sequence of each taxa. In fact, *ω* showed similar, though slightly reduced correlations, with the same empirical estimates of gene expression described above (Figure 2) This would give the impression that the same conclusions could have been gleaned using a much simpler model, both in terms of the number of parameters and the assumptions made. However, as we discussed earlier, not only is this model greatly restricted in terms of its biological feasibility, SelAC clearly performs better in terms of its fit to the data and biological realism.

**FIG. 2.**
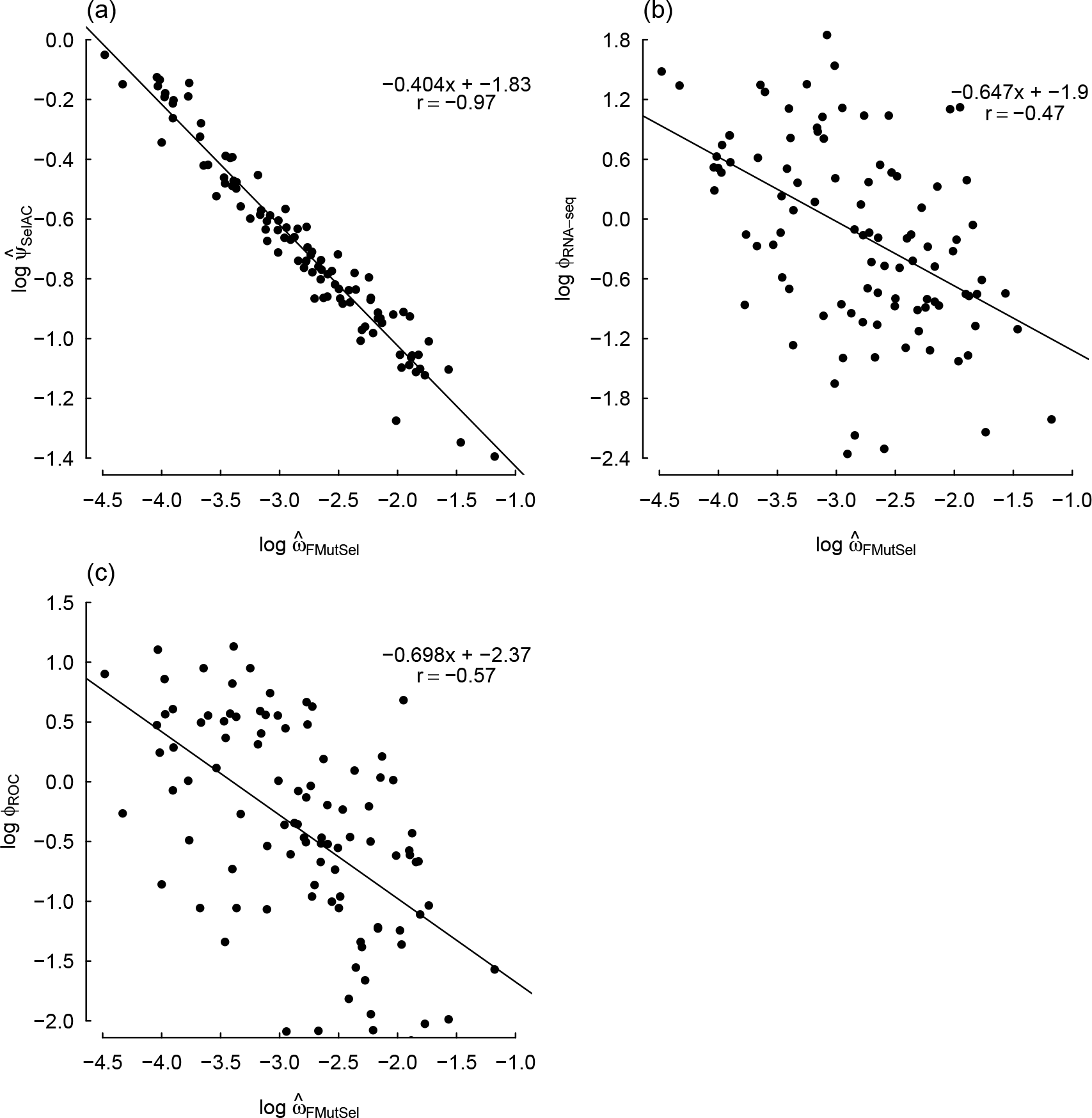
Comparisons between *ω*_FMutSel_, which is the nonsynonymous/synonymous mutation ratio in FMutSel, SelAC+Γ estimates of protein functionality production rates 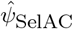 (a), RNA-Seq based measurements of mRNA abundance *ϕ*_RNA-seq_ (b), and ROC-SEMPPER’s estimates of protein translation rates *ϕ*_ROC_, which are based solely on *S. cerevisiae*’s patterns of codon usage bias (c), for *S. cerevisiae* across the 100 selected genes from Salichos and Rokas (2013). As in Figure 1, the equations in the upper right hand corner of each panel provide the regression fit and correlation coefficient.

For example, when we simulated the sequence for *S. cervisieae*, starting from the ancestral sequence under both GTR + Γ and FMutSel, the functionality of the simulated sequence moves away from the observed sequence, whereas SelAC remains near the functionality of the observed sequence (Figure 3b). This is somewhat unsurprising, given that both GTR + r and FMutSel are agnostic to the functionality of the gene, but it does highlight the improvement in biological realism in amino acid sequence evolution that SelAC provides. We do note that the adequacy of the SelAC model does vary among individual taxa, and does not always match the observed functionality. For instance, our simulations of *S. castellii* gene function is consistently higher than estimated from the data (Figure 3c). We suspect this is an indication that assuming a single set of optimal amino acid across all taxa is too simplistic. However, we cannot rule out violations of SelAC’s other model assumptions such as: a single set of Grantham weights, a single *α*_*G*_, or reductions in protein functionality **B** being solely a function of physicochemical distances *d* between sites.

**FIG. 3.**
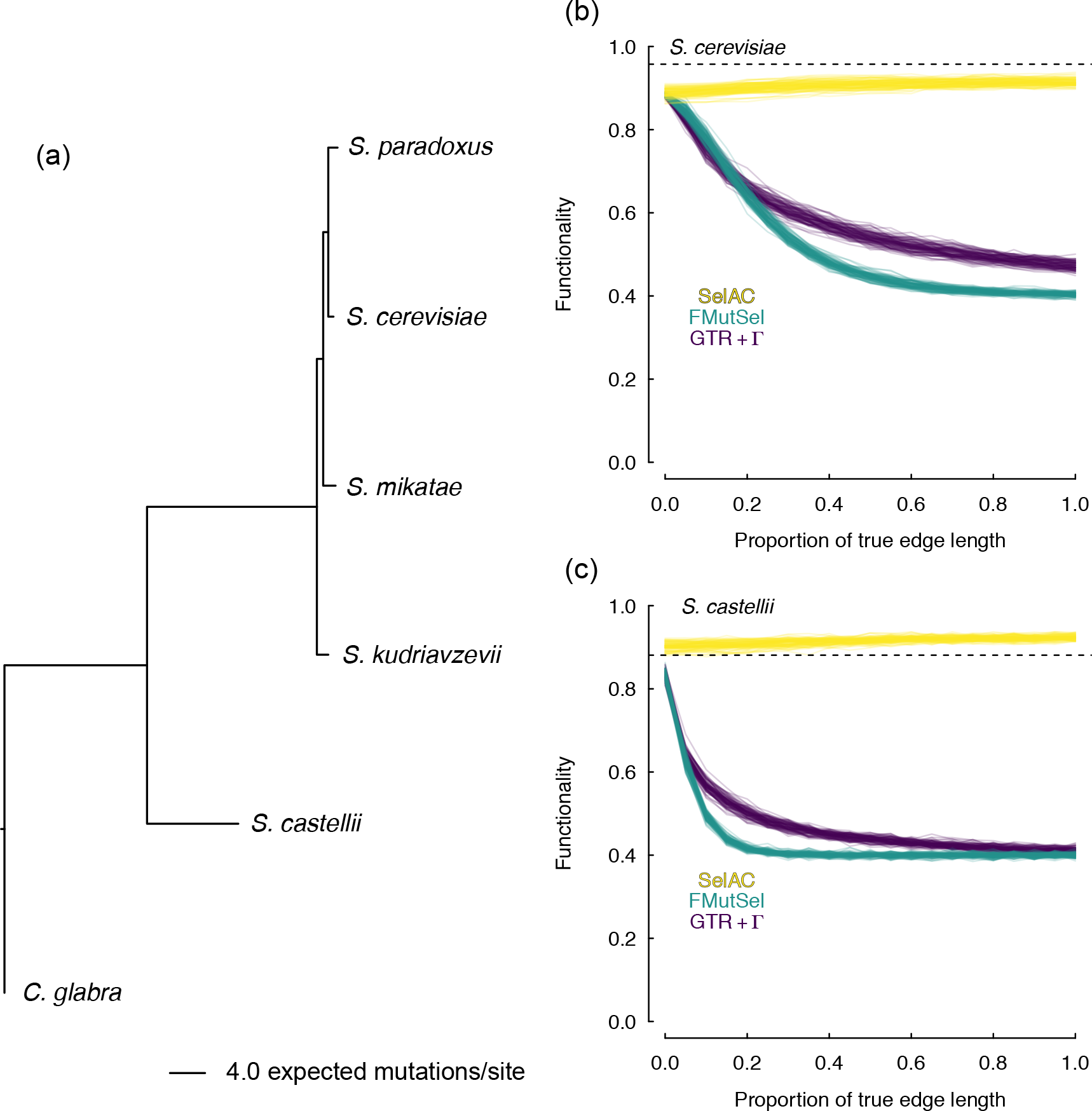
(a) Maximum likelihood estimates of branch lengths under SelAC+Γ for 100 selected genes from Salichos and Rokas (2013). Tests of model adequacy for *S. cerevisiae* (b) and *S. castellii* (c) indicated that, when these taxa are removed from the tree, and their sequences are simulated, the parameters of SelAC+Γ exhibit functionality 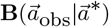 that is far closer to the observed (dashed black line) than data sets produced from parameters of either FMutSel or GTR + Γ.

## Discussion

A central goal in evolutionary biology is to quantify the nature, strength, and, ultimately, shifts in the forces of natural selection relative to genetic drift and mutation. As data set size and complexity increase, so does the amount of potential information on these forces and their dynamics. As a result, there is a need for more complex and realistic models to accomplish this goal (Goldman et al., 1996, 1998; Halpern and Bruno, 1998; Lartillot and Philippe, 2004; Thorne et al., 1996). Although extremely popular due to their elegance and computational efficiency, the utility of *ω* based models in helping us reach this goal is substantially more limited than commonly recognized. Because these *ω* models use a single substitution matrix, they are only applicable for situations in which the substitution process and shifts in the selective environment are intrinsic to the sequence, such as with positive or negative frequency dependent selection; these models do not describe stabilizing or diversifying selection as commonly envisioned (Endler, 1986; Pelmyr, 2002).

Starting with Halpern and Bruno (1998), a number of researchers have developed methods for linking site-specific selection on protein sequence and phylogenetics (e.g. Dimmic et al., 2000; Koshi and Goldstein, 2000; Koshi et al., 1999; Lartillot and Philippe, 2004; Robinson et al., 2003; Rodrigue and Lartillot, 2014; Thorne et al., 2012). Halpern and Bruno (1998) calculated a vector of 20 expected amino acid frequencies for each amino acid site, making it the most general and most parameter rich of these methods. This generality, however, comes at the cost of being purely descriptive; there is no explicit biological mechanism proposed to explain the site specific amino acid frequencies estimated. By grouping together amino sites with similar evolutionary behaviors, Lartillot and Philippe (2004) and Rodrigue and Lartillot (2014) retained the descriptive nature of Halpern and Bruno (1998) work while greatly reduced the number of model parameters needed.

SelAC follows in this tradition of using multiple substitution matrices, but includes some key advances. First, by nesting a model of a sequence’s cost-benefit function **C**/**B** within a broader model, SelAC allows us to formulate and test a hierarchical, mechanistic models of stabilizing selection. More precisely, our nested approach allows us to relax the assumption that physicochemical deviations from the optimal sequence 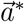 are equally disruptive at all sites within a protein. Indeed, SelAC strongly supports the hypothesis that the strength of stabilizing selection against physicochemical deviations from 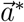 varies between sites (ΔAICc = 20,983; Table1). Second, because our substitution matrices are built on a formal description of a sequence’s cost-benefit function **C**/**B**, we are able to efficiently parameterize 20 different matrices using a relatively small number of genome-wide parameters - e.g. our physicochemical weightings, *α*_*c*_, *α*_*p*_, and *α*_*v*_, and the shape parameter *α*_*G*_ for the distribution of selective strength *G* and one gene specific expression parameter *ψ*. While the **C**/**B** function on which SelAC currently rests is very simple, nevertheless, it leads to a dramatic increase in our ability to explain the sequence data we analyzed. Importantly, because SelAC uses a formal description of a sequence’s **C**/**B**, replacing our assumptions with more sophisticated ones in the future is relatively straightforward. Third, our use of nested models also allows us to make biologically meaningful and testable predictions. By linking a gene’s expression level to the strength of purifying selection it experiences, we are able to provide coarse estimates of gene expression. This also suggests that *ω* is best explained as a proxy for gene expression, rather than the nature of selection on a sequence.

Thus, we believe our cost-benefit approach to be a substantial advance of the more simplistic *ω* models, is complementary to the work of others in the field (e.g. Rodrigue and Lartillot, 2014; Thorne et al., 2012), and, in turn, lays the foundation for more realistic work in the future. For instance, by assuming there is an optimal amino acid for each site, SelAC naturally leads to a non-symmetrical and, thus, more cogent model of protein sequence evolution. Because the strength of selection depends on an additive function of amino acid physicochemical properties, an amino acid more similar to the optimum has a higher probability of replacing a more dissimilar amino acid than the converse situation. Further, SelAC does not assume the system is always at the optimum or pessimum point of the fitness landscape, as occurs when *ω*< 1 or > 1, respectively.

Importantly, the cost-benefit approach underlying SelAC allows us to link the strength of selection on a protein sequence to its gene’s expression level. Despite its well recognized importance in determining the rate of protein evolution (e.g. Drummond et al., 2005, 2006), phylogenetic models have ignored the fact that expression levels vary between genes. In order to link gene expression and the strength of stabilizing selection on protein sequences, we simply assume that the strength of selection on a gene is proportional to the average protein synthesis rate of the gene.

One possible mechanism with some theoretical and empirical support which generates a linear relationship between the strength of selection and gene expression is the assumption of compensatory gene expression (Allison, 2012; Allison and Goulden, 2017; Brown and Elliot, 1997; King et al., 2015; Lerman et al., 2012; Thiele et al., 2012; Zanger and Schwab, 2013). That is, the assumption that any reduction in protein function is compensated for by an increase in the protein’s production rate and, in turn, abundance. For example, a mutation which reduces the functionality of the protein to 90% of the optimal protein, would require 1/0.9 = 1.11 of these suboptimal proteins to be produced relative to the optimal protein in order to maintain the same amount of that protein’s functionality in the cell. Because the energetic cost of an 11% increase in a protein’s synthesis rate is proportional to its target synthesis rate, our assumptions naturally link changes in protein functionality and changes in gene expression and its associated costs. Under what circumstances cells actually respond in this manner, remains to be determined. The fact that our method allows us to explain 13-23% of the variation in gene expression measured using RNA-Seq, suggests that this assumption is a reasonable starting point.

Furthermore, by linking expression and selection, SelAC provides a natural framework for combining information from protein coding genes with very different rates of evolution; from low expression genes providing information on shallow branches to high expression genes providing information on deep branches. This is in contrast to a more traditional approach of concatenating gene sequences together, which is equivalent to assuming the same average functionality production rate *ψ* for all of the genes, or more recent approaches where different models are fitted to different genes. Our results indicate that including a gene specific *ψ* value vastly improves SelAC fits (Table 1). Perhaps more convincingly, we find that the target functionaly production rate *ψ* and the realized average protein synthesis rate *ϕ* = *ψ*/**B** are reasonably well correlated with laboratory measurements and theoretical predictions of gene expression (Pearson *r*=0.34−0.64; Figures 1, 1, and 2). The idea that quantitative information on gene expression is embedded within intra-genomic patterns of synonymous codon usage is well accepted; our work shows that this information can also be extracted from comparative data at the amino acid level.

Of course, given the general nature of SelAC and the complexity of biological systems, other biological forces besides selection for reducing energy flux likely contribute to intergenic variation in the magnitude of stabilizing selection. Similarly, other physicochemical properties besides composition, volume, and charge likely contribute to site specific patterns of amino acid substitution. Thus, a larger and more informative set of physicochemical weights might improve our model fit and reduce the noise in our estimates of realized protein synthesis rates *ϕ*. Even if other physicochemical properties are considered, the idea of a consistent, genome wide physicochemical weighting of these terms seems highly unlikely. Since the importance of an amino acid’s physicochemical properties likely changes with its position in a folded protein, one way to incorporate such effects is to test whether the data supports multiple sets of physicochemical weights for either subsets of genes or regions within genes, rather than a single set.

Both of these points highlight the advantage of the detailed, mechanistic modeling approach underlying SelAC. Because there is a clear link between protein expression, synthesis cost, and functionality, SelAC can be extended by increasing the realism of the mapping between these terms and the coding sequences being analyzed. For example, SelAC currently assumes the optimal amino acid for any site is fixed along all branches. This assumption can be relaxed by allowing the optimal amino acid to change during the course of evolution along a branch. From a computational standpoint, the additive nature of selection between sites is desirable because it allows us to analyze sites within a gene largely independently of each other. From a biological standpoint, this additivity between sites ignores any non-linear interactions between sites, such as epistasis, or between alleles, such as dominance. Thus, our work can be considered a first step to modeling these more complex scenarios.

For example, our current implementation ignores any selection on synonymous codon usage bias (CUB) (c.f. Pouyet et al., 2016; Yang and Nielsen, 2008). Including such selection is tricky because introducing the site-specific cost effects of CUB, which is consistent with the hypothesis that codon usage affects the efficiency of protein assembly or **C**, into a model where amino acids affect protein function or **B**, results in a cost-benefit ratio **C**/**B** with epistatic interactions between all sites. These epistatic effects can likely be ignored under certain conditions or reasonably approximated based on an expectation of codon specific costs (e.g. Kubatko et al., 2016). Nevertheless, it is difficult to see how one could identify such conditions without modeling the way in which codon and amino acid usage affects **C**/**B**.

This work also points out the potential importance of further investigation into model choice in phylogenetics. For likelihood models, use of AICc has become standard. However, how one determines the appropriate number of data points in a model is more complicated than generally recognized. Common sense suggests that dataset size is increased by adding taxa and/or sites. In other words, a dataset of 1000 taxa and 100 sites must have more information on substitution models than a dataset of 4 taxa and 100 sites. Our simple analyses agree that the number of observations in a dataset (number of sites × number of taxa) should be taken as the sample size for AICc, but this conclusion likely only applies when there is sufficient independence between taxa. For instance, one could imagine a phylogeny where one taxon is sister to a polytomy of 99 taxa that have zero length terminal branches. Absent measurement error or other intraspecific variation, one would have 100 species but only two unique trait values, and the only information about the process of evolution comes from what happens on the path connecting the lone taxon to the polytomy. Although this is a rather extreme example, it seems prudent for researchers to use a simulation based approach similar to the one we take here to determine the appropriate means for calculating the effective number of data points in their data.

There are still significant shortcomings in the approach outlined here. Most worrisome are biological oversimplifications in SelAC. For example, at its heart, SelAC assumes that suboptimal proteins can be compensated for, at a cost, simply by producing more of them. However, this is likely only true for proteins reasonably close to the optimal sequence. Different enough proteins will fail to function entirely: the active site will not sufficiently match its substrates, a protein will not properly pass through a membrane, and so forth. Yet, in our model, even random sequences still permit survival, just requiring more protein production. Like the other oversimplificats previously discussed, these assumptions can be relaxed through further extension of our model.

There are also deficiencies in our implementation. Though reasonable to use for a given topology with a modest number of species, it is currently too slow for practical use for tree search. Our work serves as a proof of concept, or of utility for targeted questions where a more realistic model may be of use (placement of particular taxa, for example). Future work will encode SelAC models into a variety of mature, popular tree-search programs. SelAC also represents a challenging optimization problem: the nested models reduce parameter complexity vastly, but there are still numerous parameters to optimize, including the discrete parameter of the optimal amino acid at each site. One way to avoid the use of discrete parameters at the expense of more of them would be to have SelAC estimate the optimum physicochemical values on a per site basis rather than a specific amino acid. While this would increase the number of parameters estimated, it would have the practical advantage of continuous parameter optimization rather than discrete, and biologically would be more realistic (as it is the properties that selection “sees”, not the identity of the amino acid itself).

In spite of these difficulties, SelAC represents an important step in uniting phylogenetic and population genetic models. For example, while Dimmic et al. (2000); Koshi and Goldstein (2000); Koshi et al. (1999); Lartillot and Philippe (2004); Robinson et al. (2003); Rodrigue and Lartillot (2014); Thorne et al. (2012) are all models of constant, stabilizing selection, SelAC can be generalized further to include diversifying selection. Specifically, by letting SelAC’s sensitivity term *G*, which we now assume is ≥ 0, to take on negative values, SelAC will behave as if there is a pessimal, rather than optimal, amino acid for the given site. In this diversifying selection scenario, amino acids with physicochemical qualities more dissimilar to the pessimal amino acid are increasingly favored, potentially resulting in multiple fitness peaks.

Because SelAC infers the optimal amino acid for each site, it is substantially more parameter rich than more commonly used models such as GTR+Γ, GY94, and FMutSel. Despite this increase in number of model parameters, SelAC drastically outperforms these models with AICc values on the order of 10,000s to 100,000s. We predict that SelAC’s performance could be improved even further if we use a hierarchical approach where the optimal amino acid is not estimated on a per site basis, but rather as a vector of probability an amino acid is optimal at the gene level.

This ability to extend our model and, in turn, sharpen our thinking about the nature of natural selection on amino acid sequences illustrates the value of moving from descriptive to more mechanistic models in general and phylogenetics in particular. How frequently diversifying selection of this nature occurs is an open, but addressable, question. Regardless of the frequency at which diversifying selection occurs, another question of interest to evolutionary biologists is, “How often does the optimal/pessimal amino sequence change along any given branch?” Due to its mechanistic nature, SelAC can also be extended to include changes in the optimal/pessimal sequence over a phylogeny using a hidden markov modelling approach. Extending SelAC in these ways, will allow researchers to explicitly model shifts in selection on protein sequences and, in turn, quantify their frequency and magnitude thus deepening our understanding of biological evolution.

In summary, SelAC allows biologically relevant population genetic parameters to be estimated from phylogenetic information, while also dramatically improving fit and accuracy of phylogenetic models. By explicitly modeling the optimal/pessimal sequence of a gene, SelAC can be extended to include shifts in the optimal/pessimal sequence over evolutionary time. Moreover, it demonstrates that there remains substantially more information in the coding sequences used for phylogenetic analysis than other methods can access. Given the enormous amount of efforts expended to generate sequence datasets, it makes sense for researchers to continue developing more realistic models of sequence evolution in order to extract the biological information embedded in these datasets. The cost-benefit model we develop here is just one of many possible paths of mechanistic model development.

## Materials & Methods

### Overview

We model the substitution process as a classic Wright-Fisher process which includes the forces of mutation, selection, and drift (Berg and Lässig, 2003; Fisher, 1930; Iwasa, 1988; Kimura, 1962; McCandlish and Stoltzfus, 2014; Sella and Hirsh, 2005; Wright, 1969). For simplicity, we ignore linkage effects and, as a result of this and other assumptions, sequences evolve in a site independent manner.

Because SelAC requires twenty families of 61 × 61 matrices, the number of parameters needed to implement SelAC would, without further assumptions, be extremely large (i.e. on the order of 74,420 parameters). To reduce the number of parameters needed, while still maintaining a high degree of biological realism, we construct our gene and amino acid specific substitution matrices using a submodel nested within our substitution model, similar to approaches in Gilchrist (2007); Gilchrist et al. (2015); Shah and Gilchrist (2011).

One advantage of a nested modeling framework is that it requires only a handful of genome-wide parameters such as nucleotide specific mutation rates (scaled by effective population size *N*_*e*_), amino acid side chain physicochemical weighting parameters, and a shape parameter describing the distribution of site sensitivities. In addition to these genome-wide parameters, SelAC requires a gene *g* specific functionality expression parameter *ψ*_*g*_ which describes the average rate at which the protein’s functionality is produced by the organism or a gene’s ‘average functionality production rate’ for short (for notational simplicity, we will ignore the gene specific indicator _g_, unless explicitly needed). Currently, *ψ* is fixed across the phylogeny, though relaxing this assumption is a goal of future work. The gene specific parameter *ψ* is multiplied by additional model terms to make a composite term which *ψ*′ scales the strength and efficacy of selection for the optimal amino acid sequence relative to drift (see Implementation below). In terms of the functionality of the protein encoded, we assume that for any given gene there exists an optimal amino acid sequence 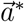 and that, by definition, a complete, error free peptide consisting of 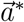 provides one unit of the gene’s functionality. We also assume that natural selection favors genotypes that are able to synthesize their proteome more efficiently than their competitors and that each savings of an high energy phosphate bond per unit time leads to a constant proportional gain in fitness *A*_0_. SelAC also requires the specification (as part of parameter optimization) of an optimal amino acid *a*\* at each position within a coding sequence. This requirement of one *a*\* per site makes our 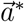 the largest category of parameters SelAC estimates. Despite the need to specify *a*\* for each site, because we use a submodel to derive our substitution matrices, SelAC estimates a relatively small number of the parameters when compared to more general approaches where the fitness of each amino acid is allowed to vary freely of any physicochemical properties (Halpern and Bruno, 1998; Lartillot and Philippe, 2004; Rodrigue and Lartillot, 2014).

As with other phylogenetic methods, SelAC generates estimates of branch lengths and nucleotide specific mutation rates. In addition, the method can also be used to make quantitative inferences on the optimal amino acid sequence of a given protein as well as the realized average synthesis rate of each protein used in the analysis. The mechanistic basis of SelAC also means it can be easily extended to include more biological realism and test more explicit hypotheses about sequence evolution.

### Mutation Rate Matrix *μ*

We begin with a 4×4 nucleotide mutation matrix ***μ*** that describes mutation rates between different bases and, in turn, different codons. For our purposes, we rely on the general unrestricted model (UNREST from Yang, 1994) because it imposes no constraints on the instantaneous rate of change between any pair of nucleotides. More constrained models, such as the Jukes-Cantor (JC), Hasegawa-Kishino-Yano (HKY), or the general time-reversible model (GTR), could also be used.

The 12 parameter UNREST model defines the relative rates of change between a pair of nucleotides. Thus, we arbitrarily set the G→T mutation rate to 1, resulting in 11 free mutation rate parameters in the 4×4 mutation nucleotide mutation matrix. The nucleotide mutation matrix is also scaled by a diagonal matrix **π** whose entries, π_*i*,*i*_, correspond to the equilibrium frequencies of each base. These equilibrium nucleotide frequencies are determined by analytically solving **π** × **Q** = 0. We use this **Q** to populate a 61 × 61 codon mutation matrix ***μ*** whose entries *μ*_*i*,*j*_ *i* ≠ *j* describes the mutation rate from codon *i* to *j* and *μ*_*i*,*j*_ = −∑_*j*_*μ*_*i*,*j*_. We generate this matrix using a “weak mutation” assumption, such that evolution is mutation limited, codon substitutions only occur one nucleotide at a time. As a result, the rate of change between any pair of codons that differ by more than one nucleotide is zero.

While the overall model does not assume equilibrium, we still need to scale our mutation matrices *μ* by a scaling factor *S*. As traditionally done, we rescale our time units such that at equilibrium, one unit of branch length represents one expected mutation per site (which equals the substitution rate under neutrality). More explicitly, *S* = −(∑_*i*∈codons_*μ*_*i*,*i*_*π*_*i*,*i*_) where the final mutation rate matrix is the original mutation rate matrix multiplied by 1/*S*.

### Protein Synthesis Cost-Benefit Function *η*

SelAC links fitness to the product of the cost-benefit function of a gene *η* and the organism’s average target synthesis rate of the functionality provided by gene *ψ*. As a result, the average flux energy an organism spends to meet its target functionality provided by the gene is *η* × *ψ*. Compensatory changes that allow an organism to maintain functionality even with loss of one or both copies of a gene are widespread. There is evidence of compensation for protein function. Metabolism with gene expression models (ME-models) link those factors to successfully make predictions about response to perturbations in a cell (King et al., 2015; Lerman et al., 2012). For example, an ME-model for *E. coli* successfully predicted gene expression levels in vivo (Thiele et al., 2012). Here we assume that for finer scale problems than entire loss (for example, a 10% loss of functionality) the compensation is more production of the protein. The particular type of dosage compansation assumed by SelAC in respondse to stress (e.g. reduced functionality) is commonly assumed in microbial ecology (Allison, 2012; Allison and Goulden, 2017). Our assumption is also consistent with the Michaelis-Menten enzyme kinetics. Moreover, there is evidence that mutations can influence expression level, though this does not always match our expression compensation assumption (Brown and Elliot, 1997; Zanger and Schwab, 2013). In order to link genotype to our cost-benefit function *η* = **C**/**B**, we begin by defining our benefit function **B**.

#### Benefit

Our benefit function **B** measures the functionality of the amino acid sequence 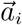 encoded by a set of codons 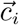, i.e. 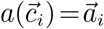 relative to that of an optimal sequence 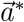. By definition, 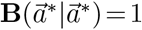 and 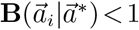 for all other sequences. We assume all amino acids within the sequence contribute to protein function and that this contribution declines as an inverse function of physicochemical distance from each amino acid to the optimal one. Formally, we assume that

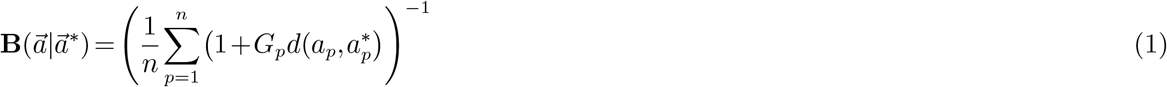

where *n* is the length of the protein, 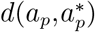 is a weighted physicochemical distance between the amino acid encoded at a given position *p* and 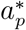 is the optimal amino acid for that position. There are many possible measures for physiochemical distance; we use Grantham (1974) distances by default, though others may be chosen. For simplicity, we assume all nonsense mutations are lethal by defining the the physicochemical distance between a stop codon and a sense codon as ∞. The term *G*_*p*_ describes the sensitivity of the protein’s function to physicochemical deviation from the optimimum at site position *p*. We assume that *G*_*p*_ ~Gamma(shape = *α*_*G*_,rate = *α*_*G*_) in order to ensure 𝔼(*G*_*p*_) = 1. Given the definition of the Gamma distribution, the variance in *G*_*p*_ is equal to shape/rate^2^ = 1/*α*_*G*_. We note that at the limit of *α*_*G*_ → ∞ to, the model becomes equivalent to assuming uniform site sensitivity where *G*_*p*_ = 1 for all positions *p*. Further, 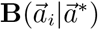 is inversely proportional to the average physicochemical deviation of an amino acid sequence 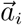 from the optimal sequence 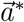 weighted by each site’s sensitivity to this deviation. 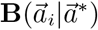 can be generalized to include second and higher order terms of the distance measure *d*.

#### Cost

Protein synthesis involves both direct and indirect assembly costs. Direct costs consist of the high energy phosphate bonds ~ *P* of ATPs or GTPs used to assemble the ribosome on the mRNA, charge tRNA’s for elongation, move the ribosome forward along the transcript, and terminate protein synthesis. As a result, direct protein assembly costs are the same for all proteins of the same length. Indirect costs of protein assembly are potentially numerous and could include the cost of amino acid synthesis as well the cost and efficiency with which the protein assembly infrastructure such as ribosomes, aminoacyl-tRNA synthetases, tRNAs, and mRNAs are used. When these indirect costs are combined with sequence specific benefits, the probability of a mutant allele fixing is no longer independent of the rest of the sequence (Gilchrist et al., 2015) and, as a result, model fitting becomes substantially more complex. Thus for simplicity, in this study we ignore indirect costs of protein assembly that vary between genotypes and define,

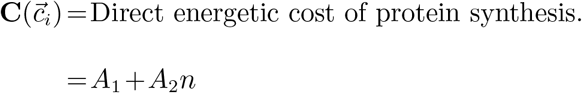

where, *A*_1_ and *A*_2_ represent the direct cost, in high energy phosphate bonds, of ribosome initiation and peptide elongation, respectively, where *A*_1_ = *A*_2_=4 ~ *P*.

### Defining Physicochemical Distances

Assuming that functionality declines with an amino acid *a*_*i*_’s physicochemical distance from the optimum amino acid *a*\* at each site provides a biologically defensible way of mapping genotype to protein function that requires relatively few free parameters. In addition, SelAC naturally lends itself to model selection since one could compare the quality of SelAC fits using different mixtures of physicochemical properties. Following (Grantham, 1974), we focus on using composition *c*, polarity *p*, and molecular volume *v* of each amino acid’s side chain residue to define our distance function, but the model and its implementation can flexibly handle a variety of properties. We use the Euclidian distance between residue properties where each property *c*, *p*, and *v* has its own weighting term, *α*_*c*_, *α*_*p*_, *α*_*v*_, respectively, which we refer to as ‘Grantham weights’. Because physicochemical distance is ultimately weighted by a gene’s specific average protein synthesis rate *ψ*, another parameter we estimate, there is a problem with parameter identifiablity. The scale of gene expression is affected by how we measure physicochemical distances which, in turn, is determined by our choice of Grantham weights. As a result, by default we set *α*_*v*_=3.990 × 10^−4^, the value originally estimated by Grantham, and recognize that our estimates of *α*_*c*_ and *α*_*p*_ and *ψ* are scaled relative to this choice for *α*_*v*_. More specifically,

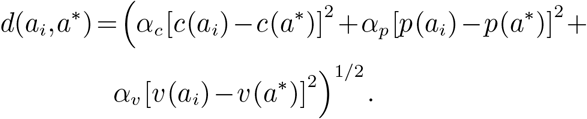

### Linking Protein Synthesis to Allele Substitution

Next we link the protein synthesis cost-benefit function *η* of an allele with its fixation probability. First, we assume that each protein encoded within a genome provides some beneficial function and that the organism needs that functionality to be produced at a target average rate *ψ*. Again, by definition, the optimal amino acid sequence for a given gene, 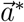, produces one unit of functionality, i.e. 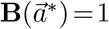. Second, we assume that the actual average rate a protein is synthesized *ϕ* is regulated by the organism to ensure that functionality is produced at rate *ψ*. As a result, it follows that 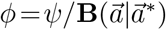 and the energetic burden of a suboptimal amino acid increases the more it decreases the protein’s functionality, **B**. In other words, the average production rate of a protein 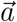 with relative functionality 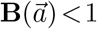 must be 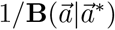 times higher than the production rate needed if the optimal amino acid sequence 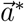 was encoded since 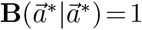. For example, a cell with an allele 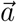 where 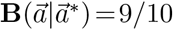 would have to produce the protein at rate *ϕ*=10/9×*ψ*=1.11*ψ*. Similarly, a cell with an allele 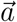 where 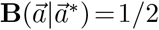 will have to produce the protein at *ϕ*=2*ψ*. In contrast, a cell with the optimal allele 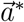 would have to produce the protein at rate *ϕ*=*ψ*.

Third, we assume that every additional high energy phosphate bond, ~*P*, spent per unit time to meet the organism’s target function synthesis rate *ψ* leads to a slight and proportional decrease in fitness *W*. This assumption, in turn, implies

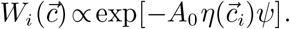

where *A*_0_, again, describes the proportional decline in fitness with every ~*P* wasted per unit time. Because *A*_0_ shares the same time units as *ψ* and *ϕ* and only occurs in SelAC in conjunction with *ψ*, we do not need to explicitly identify our time units. Instead, we recognize that our estimates of *ψ* share an unknown scaling term.

Correspondingly, the ratio of fitness between two genotypes is,

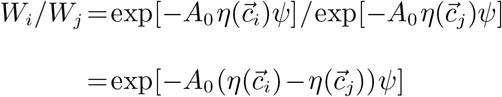

Given our formulations of **C** and **B**, the fitness effects between sites are multiplicative and, therefore, the substitution of an amino acid at one site can be modeled independently of the amino acids at the other sites within the coding sequence. As a result, the fitness ratio for two genotypes differing at multiple sites simplifies to

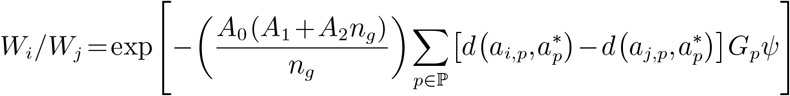

where ℙ represents the codon positions in which 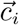 and 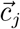 differ. Fourth, we make a weak mutation assumption, such that alleles can differ at only one position at any given time, i.e. |ℙ| = 1, and that the population is evolving according to a Wright-Fisher process. As a result, the probability a new mutant, *j*, introduced via mutation into a resident population *i* with effective size *N*_*e*_ will go to fixation is,

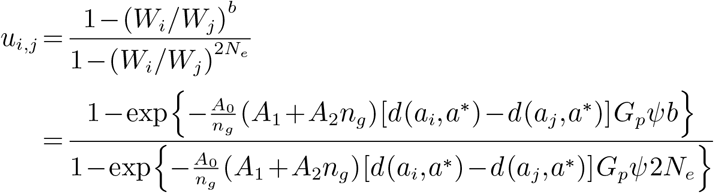

where *b* = 1 for a diploid population and 2 for a haploid population (Berg and Lässig, 2003; Iwasa, 1988; Kimura, 1962; Sella and Hirsh, 2005; Wright, 1969). Finally, assuming a constant mutation rate between alleles *i* and *j*, *μ*_*i*,*j*_, the substitution rate from allele *i* to *j* can be modeled as,

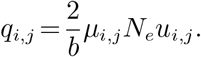

where, given the substitution model’s weak mutation assumption, *N*_*e*_μ≪1. In the end, each optimal amino acid has a separate 61 × 61 substitution rate matrix **Q**_*a*_, which incorporates selection for the amino acid (and the fixation rate matrix this creates) as well as the common mutation parameters across optimal amino acids. This results in the creation of 20 **Q** matrices, one for each amino acid and each with 3,721 entries which are based on a relatively small number of model parameters (one to 11 mutation rates, two free Grantham weights, the cost of protein assembly, *A*_1_ and *A*_2_, the gene specific target functionality synthesis rate *ψ*, and optimal amino acid at each position *p*, 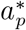). These model parameters can either be specified *a priori* and/or estimated from the data.

Given our assumption of independent evolution among sites, it follows that the probability of the whole data set is the product of the probabilities of observing the data at each individual site. Thus, the likelihood 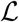 of amino acid *a* being optimal at a given site position *p* is calculated as

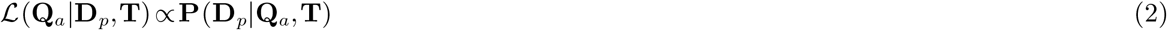

In this case, the data, **D**_*p*_, are the observed codon states at position *p* for the tips of the phylogenetic tree with topology **T**. For our purposes we take **T** as given, but it could be estimated as well. The pruning algorithm of Felsenstein (1981) is used to calculate 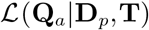. The log of the likelihood is maximized by estimating the genome scale parameters which consist of 11 mutation parameters, which are implicitly scaled by 2*N*_*e*_/*b*, and two Grantham distance parameters, *α*_*c*_ and *α*_*p*_, and the sensitivity distribution parameter *α*_*G*_. Because *A*_0_ and *ψ*_*g*_ always co-occur and are scaled by *N*_*e*_, for each gene *g* we estimate a composite term 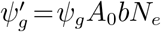 and the optimal amino acid for each position 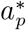 of the protein. When estimating *α*_*G*_, the likelihood then becomes the average likelihood which we calculate using the generalized Laguerre quadrature with *k* = 4 points (Felsenstein, 2001).

Finally, we note that because we infer the ancestral state of the system, our approach does not rely on any assumptions of model stationarity. Nevertheless, as our branch lengths grow the probability of observing a particular amino acid *a* at a given site approaches a stationary value proportional to 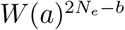 and any effects of mutation bias (Sella and Hirsh, 2005).

### Implementation

All methods described above are implemented in the new R package, selac available through GitHub (https://github.com/bomeara/selac) which will be uploaded to CRAN once peer review has completed. Our package requires as input a set of fasta files that each contain an alignment of coding sequence for a set of taxa, and the phylogeny depicting the hypothesized relationships among them. In addition to the SelAC models, we implemented the GY94 codon model of Goldman and Yang (1994), the FMutSel mutation-selection model of Yang and Nielsen (2008), and the standard general time-reversible nucleotide model that allows for r distributed rates across sites. These likelihood-based models represent a sample of the types of popular models often fit to codon data.

For the SelAC models, the starting guess for the optimal amino acid at a site comes from ‘majority’ rule, where the initial optimum is the most frequently observed amino acid at a given site (ties resolved randomly). Our optimization routine utilizes a four stage hill climbing approach. More specifically, within each stage a block of parameters are optimized while the remaining parameters are held constant. The first stage optimizes the block of branch length parameters. The second stage optimizes the block of gene specific composite parameters 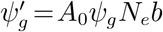. The third stage optimizes SelAC’s parameters shared across the genome *α*_*c*_ and *α*_*p*_, and the sensitivity distribution parameter *α*_*G*_. The fourth stage estimates the optimal amino acid at each site *a*\*. This entire four stage cycle is repeated six more times, using the estimates from the previous cycle as the initial conditions for the new one. The search is terminated when the improvement in the log-likelihood between cycles is less than 10^−8^ at which point we consider the ML solution found and the search is terminated. For optimization of a given set of parameters, we rely on a bounded subplex routine (Rowan, 1990) in the package `NLoptR` (Johnson, 2012) to maximize the log-likelihood function. To ensure the robustness of our results, we perform a set of independent analyses with different sets of naive starting points with respect to the gene specific composite *ψ*′ parameters, *α*_*c*_, and *α*_*p*_ and were able to repeatedly reach the same log-likelihood (lnL) peak in our parameter space. Confidence in the parameter estimates can be generated by an ‘adaptive search’ procedure that we implemented to provide an estimate of the parameter space that is some pre-defined likelihood distance (e.g., 2 lnL units) from the maximum likelihood estimate (MLE), which follows Beaulieu and O’Meara (2016) and Edwards (1984).

We note that our current implementation of SelAC is painfully slow, and is best suited for data sets with relatively few number of taxa (i.e. < 10). This limitation is largely due to the size and quantity of matrices we create and manipulate to calculate the log-likelihood of an individual site. Ongoing work will address the need for speed, with the eventual goal of implementing SelAC in popular phylogenetic inference toolkits, such as RevBayes (Hhna et al., 2016), PAML (Yang, 2007) and RAxML (Stamatakis, 2006).

### Simulations

We evaluated the performance of our codon model by simulating datasets and estimating the bias of the inferred model parameters from these data. Our ‘known’ parameters under a given generating model were based on fitting SelAC to the 106 gene data set and phylogeny of Rokas et al. (2003). The tree used in these analyses is outdated with respect to the current hypothesis of relationships within *Saccharomyces*, but we rely on it simply as a training set that is separate from our empirical analyses (see section below). Bias in the model parameters were assessed under two generating models: one where we assumed a model of SelAC assuming uniform sensitivity across sites (i.e. *G*_*p*_ = 1 for all sites, i.e. *α*_*G*_ = ∞), and one where we used the Gamma distribution joint shape and rate parameter *α*_*G*_ estimated from the empirical data. Under each of these two scenarios, we used parameter estimates from the corresponding empirical analysis and simulated 50 five-gene data sets. For the gene specific composite parameter 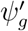 the ‘known’ values used for the simulation were five evenly spaced points along the rank order of the estimates across the 106 genes. The MLE estimate for a given replicate were taken as the fit with the highest log-likelihood after running five independent analyses with different sets of naive starting points with respect to the composite 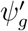 parameter, *α*_*c*_, and *α*_*p*_. All analyses were carried out in our `selac` R package.

### Analysis of yeast genomes & tests of model adequacy

We focus our empirical analyses on the large yeast data set and phylogeny of Salichos and Rokas (2013). As a model system, the yeast genome is an ideal system to examine our phylogenetic estimates of gene expression and its connection to real world measurements of these data within individual taxa. The complete data set of Salichos and Rokas (2013) contain 1070 orthologs, where we selected 100 at random for our analyses. We also focus our analyses on *Saccharomyces sensu stricto* and their sister taxon *Candida glabrata*, and we used the phylogeny depicted in Fig. 1 of Salichos and Rokas (2013) for our fixed tree. We fit the two SelAC models described above (i.e., SelAC and SelAC+Γ), as well as two codon models, GY94 and FMutSel, and a standard GTR + Γ nucleotide model. The FMutSel model assumes that the amino acid frequencies are determined by functional requirements of the protein while the other models make no assumptions about amino acid frequencies. In all cases, we assumed that the model was partitioned by gene, but with branch lengths linked across genes.

For SelAC, we compared our estimates of *ϕ*′=*ψ*′/**B**, which represents the average protein synthesis rate of a gene, to estimates of gene expression from empirical data. Specifically, we examined gene expression data for five of the six species measured during log-growth phase. Gene expression in this context corresponds to mRNA abundances, which were measured using either microarrays (*C. glabrata* and *S. castellii*, or RNA-Seq (*S. paradoxus*, *S. mikatae*, and *S. cerevisiae*). We obtained expression data for the remaining species, *S. kudriavzevii*, which was measured at the beginning of the stationary phase from the Gene Expression Omnibus (GEO). Saccharomyces, however, only enter the stationary growth phase in response to severe stress, such as starvation. In addition, only 56 % of the genes examined with SelAC had expression measurements available. For these reasons, we excluded *S. kudriavzevii* from our comparisons of empirical gene expression.

For further comparison, we also predicted the average protein synthesis rate for each gene *ϕ* by analyzing gene and genome-wide patterns of synonymous codon usage using ROC-SEMPPR (Gilchrist et al., 2015) for each individual genome. While, like SelAC, ROC-SEMPPR uses codon level information, it does not rely on any interspecific comparisons and, unlike SelAC, uses only the intra- and inter-genic frequencies of synonymous codon usage as its data. Nevertheless, ROC-SEMPPR predictions of gene expression *ϕ* correlates strongly (Pearson *r*=0.53−0.74) with a wide range of laboratory measurements of gene expression (Gilchrist et al., 2015).

While one of our main objectives was to determine the improvement of fit that SelAC has with respect to other standard phylogenetic models, we also evaluated the adequacy of SelAC. Model fit, measured with assessments such as the Akaike Information Criterion (AIC), can tell which model is least bad as an approximation for the data, but it does not reveal whether a model is actually doing a good job of representing the data. An adequate model does the latter, one measure of which is that data generated under the model resemble real data (Goldman, 1993). For example, Beaulieu et al. (2013) assessed whether parsimony scores and the size of monomorphic clades of empirical data were within the distributions of simulated data under a new model and the best standard model; if the empirical summaries were outside the range for each, it would have suggested that neither model was adequately modeling this part of the biology.

In order to test adequacy for a given gene we first remove a particular taxon from the data set and the phylogeny. A marginal reconstruction of the likeliest sequence across all remaining nodes is conducted under the model, including the node where the pruned taxon attached to the tree. The marginal probabilities of each site are used to sample and assemble the starting coding sequence. This sequence is then evolved along the branch, periodically being sampled and its current functionality assessed. We repeat this process 100 times and compare the distribution of trajectories against the observed functionality calculated for the gene. For comparison, we also conducted the same test, by simulating the sequence under the standard GTR + Γ nucleotide model, which is often used on these data but does not account for the fact that the sequences are protein coding, and under FMutSel, which includes selection on codons but in a fundamentally different way as our model.

### The appropriate estimator of bias for AIC

As part of the model set described above, we also included a reduced form of each of the two SelAC models, SelAC and SelAC+Γ. Specifically, rather than optimizing the amino acid at any given site, we assume the the most frequently observed amino acid at each site is the optimal amino acid *a*\*. We refer to these ‘majority rule’ models as SelAC_*M*_ and SelAC_*M*_+Γ and note that these majority rule formulations greatly accelerate model fitting.

Since these majority rule models assume that the optimal amino acids are known prior to fitting of our model, it is tempting to reduce the count of estimated parameters in the model by the number of parameters estimated using majority rule. While using majority rule does not necessarily provide the most likely parameter estimate, it nevertheless uses the data to generate the estimate and, represents a parameter estimated from the data. Thus, despite having become standard behavior in the field of phylogenetics, this reduction is statistically inappropriate. Because the difference in the number of parameters *K* when counting or not counting the number of nucleotide sites drops out when comparing nucleotide models with AIC, this statistical issue does not apply to nucleotide models. It does, however, matter for AICc, where *K* and the sample size *n* combine in the penalty term. This also matters in our case, where the number of estimated parameters for the majority rule estimation differs based on whether one is looking at codons or single nucleotides.

In phylogenetics two variants of AICc are used. In comparative methods (e.g. Beaulieu et al., 2013; Butler and King, 2004; O’Meara et al., 2006) the number of data points, *n*, is taken as the number of taxa. More taxa allow the fitting of more complex models, given more data. However, in DNA evolution, which is effectively the same as a discrete character model used in comparative methods, the *n* is taken as the number of sites. Obviously, both cannot be correct. This uncertainty was highlighted by Posada and Buckley (2004): they chose to use number of sites, but mentioned in their discussion that sample size also depends on the number of taxa. Sullivan and Joyce (2005) also mention that while the number of sites is often taken as sample size, whether that is appropriate in phylogenetics is not entirely clear. One approach incorporating both number of taxa and sites in calculating AICc is the program SURFACE implemented by Ingram and Mahler (2013), which uses multiple characters and taxa. While its default is to use AIC to compare models, if one chooses to use AICc, the number of samples is taken as the product of number of sites and number of taxa.

Recently, Jhwueng et al. (2014) performed an analysis that investigated what variant of AIC and AICc worked best as an estimator, but the results were inconclusive. Here, we have adopted and extended the simulation approach of Jhwueng et al. (2014) in order to examine a large set of different penalty functions and how well they approximate the remaining portion of the Kullback-Liebler (KL) divergence between two models after accounting for the deviance (i.e., 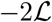) (see Appendix 1 for more details).

## Acknowledgements

This work was supported in part by NSF Awards MCB-1120370 (MAG and RZ) and DEB-1355033 (BCO, MAG, and RZ) with additional support from The University of Tennessee Knoxville and University of Arkansas (JMB). JJC and JMB received support as Postdoctoral Fellows and CL received support as a Graduate Student Fellow at the National Institute for Mathematical and Biological Synthesis, an Institute sponsored by the National Science Foundation through NSF Award DBI-1300426, with additional support from UTK. The authors would like to thank Premal Shah, Todd Oakley, and two anonymous reviewers for their helpful criticisms and suggestions for this work.

